# Transfer learning with deep convolutional neural networks for classifying cellular morphological changes

**DOI:** 10.1101/345728

**Authors:** Alexander Kensert, Philip J Harrison, Ola Spjuth

## Abstract

Quantification and identification of cellular phenotypes from high content microscopy images have proven to be very useful for understanding biological activity in response to different drug treatments. The traditional approach has been to use classical image analysis to quantify changes in cell morphology, which requires several non-trivial and independent analysis steps. Recently convolutional neural networks have emerged as a compelling alternative, offering good predictive performance and the possibility to replace traditional workflows with a single network architecture. In this study we applied the pre-trained deep convolutional neural networks ResNet50, InceptionV3 and InceptionResnetV2 to predict cell mechanisms of action in response to chemical perturbations for two cell profiling datasets from the Broad Bioimage Benchmark Collection. These networks were pre-trained on ImageNet enabling much quicker model training. We obtain higher predictive accuracy than previously reported, between 95 and 97% based on “leave-one-compound-out” cross-validation. The ability to quickly and accurately distinguish between different cell morphologies from a scarce amount of labelled data illustrates the combined benefit of transfer learning and deep convolutional neural networks for interrogating cell-based images.

## 1 Introduction

High-content screening (HCS) has proven to be a useful and successful technique to identify and quantify cell phenotypes [1, 2]. Although conventional approaches for classification of phenotypes using cell-images have shown positive results [3, 4, 5, 6, 7], they require several non-trivial data analysis steps. An example is Ljosa et al. [7] and their pipeline workflows which include cellular segmentation, feature extraction, profiling methods (e.g. factor analysis) and a nearest neighbour classifier. Cell segmentation algorithms typically require manual adjustments for each new experimental setup [4] and feature extraction tends to rely on “hand-crafted” features, such as those related to texture and shape (several of which are computationally expensive to measure). PCA, assuming a linear mapping, is then often used to reduce the dimensionality of these high dimensional (> 500) and highly correlated feature sets [8].

Convolutional neural networks (CNNs) have recently brought about breakthroughs in computer vision and image processing — CNNs automatically discover the features needed for classification of images based solely on the raw pixel intensity data [9]. This supervised feature learning technique has shown to be superior to using traditional hand-crafted features [10, 11], and the combination of segmentation and classification in a single framework [12] means that image classification can be performed without the need for prior cell segmentation (a complex task that often requires careful consideration and significant computation [13]). A recent survey shows a rapid growth in the application of deep learning to medical image analysis [14], with several studies outperforming medical expert classification. A convenient property of CNNs is that the pipeline workflow of the traditional methods are taken care of by the network itself; and by convolving filters (the weights/parameters of the network) on input layers, local connectivity and parameter sharing keeps the number of parameters relatively low, even for a deeper network.

A major bottleneck when applying supervised CNNs to cell images is the scarcity of labelled data. Importantly, studies have shown that reusing models trained on different tasks reduced these problems [15, 16]. Josinski et al. [17] note that transferability of features depends on the distance between the base task and the target task. However, the features from distant tasks may still perform better than random features. The study also illustrated that initializing the network with pre-trained features improved the generalization even after considerable fine-tuning to the target dataset. Further, Zhang et al. [18] showed that features trained on natural images (ImageNet [19]) could be transferred to biological data. Neslihan et al. [20] used pre-trained models on natural images and facial images for cell nucleus classification where the performance of transfer learning and learning from scratch were compared. All of their pretrained models had better predictive performance whilst requiring less training time. Phan et al. [21] also successfully utilized transfer learning on bioimages and outperformed all other methods on the mitosis detection dataset of the ICPR2012 contest. The utility of transfer learning is in part due to the fact that the initial CNN layers capture low-level features, like edges and blobs — characteristics commonly shared between different types of images.

The bbbc (broad bioimage benchmark collection) is an important publicly available collection of microscopy images intended for validating image-analysis algorithms [22]. Various algorithms have been tested and validated on these datasets — ranging from traditional pipeline workflows to deep learning techniques [5, 7, 23, 24, 25]. Pawlowski et al. [24] utilized transfer learning without fine-tuning to extract features, and Ando et al. [23] used a pre-trained model on consumer images and further transformation techniques to attain the top accuracy on this benchmark dataset of 96%. However, research on transfer learning and fine tuning of CNNs on these bbbc datasets is scarce — it is therefore important to investigate and compare this technique with the existing analysis tools that have been applied to the bbbc datasets.

In this study we present state-of-the-art deep convolutional neural networks pre-trained on natural images, with minor image pre-processing and without segmentation. These models were used to predict mechanisms of action (MoA) and nucleus translocation, based only on pixel intensities which automatically pass through the network to give the final predictions. We used two different bbbc datasets: bbbc021v1 and bbbc014v1 [22], to evaluate the models predictive performance as well as to visualize the filter output activations (feature maps) throughout the network. This visualization was done to understand the different levels of abstraction processed and the transferability of the networks. After the parameter values were transferred the networks were fine-tuned to fit the data — the transferred parameters can thus be thought of as good initial parameter values. Although no comparison with randomized initialization of parameter values was done, we hypothesized that the pre-trained parameters would improve performance both in terms of accuracy and learning time.

## 2 Method

### 2.1 Data

#### 2.1.1 Datasets

The first dataset, bbbc021v1, used in this study contains MCF-7 breast cancer images available from the Broad Bioimage Benchmark Collection [22] — wells were fixed, labeled for DNA, F-actin, B-tubulin, and imaged by fluorescence microscopy as described by Caie et al [26]. We used 38 out of 113 compounds (1-7 concentrations each) which were annotated with one MoA, resulting in a total of 103 treatments (compound-concentration pairs) and 12 different MoAs (for more information see https://data.broadinstitute.org/bbbc/BBBC021/).

The second dataset used in this study was bbbc014v1 provided by Ilya Ravkin, and is also available from the Broad Bioimage Benchmark Collection [22]. The images are Human U2OS cells of cytoplasm to nucleus translocation of the MCF-7 and A549 (human alveolar basal epithelial) in response to TNFα concentrations. For each well there was one field with a nuclear counterstain (DAPI) and one field with a signal stain (FITC). A total of 96-wells with 12 concentration points and 4 replicate rows for each cell type (for more information see https://data.broadinstitute.org/bbbc/BBBC014/). In this study, the four highest (labelled positive) and four lowest concentrations (including 0 concentration; labelled negative) were used for each cell type.

#### 2.1.2 Image preprocessing

For the bbbc021v1 dataset, the three different 16-bit range channels (labelled for DNA, F-actin, B-tubulin) were stacked into a three channelled image. The images were then normalized plate-wise, by subtracting the mean pixel intensities of DMSO images (the control samples) and then dividing by the standard deviation of their pixel intensities. After the normalization an Anscombe transformation [27] was performed followed by a mapping to an 8-bit range.

Similarly, for the bbbc014v1 dataset, the different channels (in this case 2) of each sample were stacked with an addition of a zero matrix to create a three-channel input. These images were in 8-bit range and were variance stabilized by the Anscombe transformation, and then mapped back to an 8-bit range.

The resulting images for both datasets were cropped into 16 images (bbbc014v1) and 4 images (bbbc021v1) to increase the number of training samples.

### 2.2 CNN architectures

Three different state-of-the-art architectures were implemented in Keras [28]: Resnet50 [29], InceptionV3 [30] and InceptionResnetV2 [31]. They were all pretrained on the ImageNet dataset, containing 13 million natural images [19].

#### 2.2.1 Residual network

Utilizing a very deep CNN can have a negative effect on model performance — arising from the difficulty in finding suitable parameters for the deeper layers. Adding further layers to a suitably deep model can lead to higher training error not caused by overfitting [29, 32, 33]. Residual networks use residual mapping *H*(*x*) = *F(x)* + *x*, where *x* is the original feature vector (identity mapping) added to the deeper version of the network *F(x)* (output of the stacked layers). Importantly, if the mappings are optimal, it is easier for the network to push the residuals to zero than fitting an identity mapping with stacks of nonlinear layers [29]. The implication of this is that although F(x) is not learning anything, the output will simply be an identity mapping *x*. Thus, in the worst-case scenario the output equals the input, and in the best-case some important features are learned. Residual mappings therefore assist in avoiding the degradation problem that occurs for very deep CNNs. Another important aspect of residual networks is the intermediate normalization layers (also called batch normalization), which help to solve the problem of vanishing and exploding gradients.

The Residual network used in this study had 50 layers (49 convolutional layers and a final fully connected classification layer), based on ResNet50 from the paper “Deep Residual Learning for Image Recognition” [29].

#### 2.2.2 Inception network

It is often difficult to determine the best network filter sizes and whether or not to use pooling layers. To overcome this inception architectures use many different filter sizes and pooling layers in parallel (an inception block), the outputs of which are concatenated and inputted to the next block. In this way the network chooses which filter sizes or combinations thereof to use. To solve the problem of a large increase in computational cost, the inception networks utilize 1×1 convolutions to shrink the volume of the next layer. This network architecture was introduced by Szegedy et al. [34] to make a network deeper and wider, hence more powerful, whilst keeping the computational cost low. The Inception network can thus go very deep and, like Resnet, utilizes intermediate normalization layers to avoid vanishing and exploding gradients.

The inception network used in this study was the InceptionV3 from the paper ‘‘Rethinking the Inception Architecture for Computer Vision” [30], excluding the auxiliary classifiers. This network had 95 layers in total, a number much larger than ResNet50 due to the width of each inception block.

#### 2.2.3 Inception-residual network

Szegedy et al. [31] evaluated a network combining inception blocks and residuals (similar to the ResNet50 residuals). They showed an improvement in training speed after introducing these residuals, making it possible to implement even deeper networks at a reasonable cost.

In this study, we implemented an inception-resnet architecture based on the InceptionResnetV2 from the paper “Inception-v4, Inception-ResNet and the Impact of Residual Connections on Learning” [31]. This network is even than ResNet50 and InceptionV3 combined — totalling 245 layers.

### 2.3 Downsampling and data augmentation

Before the bbbc021v1 images were inputted into the network they were down-sampled to have the same dimensions as the images used for the pretrained network: 224×224×3 for ResNet50 and 299×299×3 for InceptionV3 and InceptionResnetV2. For the bbbc014v1 dataset all images were down-sampled to have dimensions of 256×256×3. To increase the number of training examples for bbbc021v1, the input images were randomly rotated and mirrored. Further, jitter, blur and Gaussian noise were then randomly applied to both prevent the network from identifying noise as important features and to augment the data further.

### 2.4 Model evaluation and deep visualization

#### 2.4.1 Model evaluation

To evaluate the models of the bbbc021v1 dataset we used a “leave-one-compound-out” cross-validation — resulting in a 38-fold cross-validation. In each fold predictions were made for all the treatments of the excluded compound. An element-wise median over the replicates was first calculated to obtain a prediction vector for each well. These vectors were then used to calculate the element-wise median over the wells, to obtain prediction vectors for each treatment. Finally, the highest values in the resulting 12-dimensional prediction vectors, containing the MoA predictions for the treatments, decided the models final predictions for the treatments. This procedure was repeated for all cross-validation folds, resulting in a total of 103 final predictions.

For the bbbc014v1 dataset we used a 2-fold cross-validation where one cell-line was “left-out” as the test set, while the other was used for training. At test time, a prediction was made for each image (well) — resulting in 32 predictions for each fold.

#### 2.4.2 Activation maximization

To compare the pre-trained models and fine-tuned models (fit to our MoA data), we compared a pre-trained ImageNet ResNet50 model and a fine-tuned ResNet50 model (trained on all images for ten epochs) and contrasted a selection of their filters. We used the high-level Keras visualization toolkit *keras-vis* [35] to do this and applied an activation maximization function to generate an input image that maximizes certain filter output activations. This allows us to understand the input patterns that activate certain filters. The network filters of the deeper layers learn more abstract representations of the data, which are reflected in their more intricately structured visualizations [36].

## 3 Results and Discussion

To evaluate the models on the bbbc021v1 dataset we predicted the MoA for each treatment of the left-out compound for each fold in the cross-validation - hence testing our deep CNN models on unseen compounds and its treatments 103 times. We illustrate the accuracies of these predictions by plotting confusion matrices for all MoAs (Figure 1). ResNet50, InceptionV3 and InceptionResnetV2 attained mean accuracies of 97%, 97% and 95% respectively — thus comparing well with previous state-of-the-art algorithms in terms of hard predictions, where our ResNet50 and InceptionV3 applications reached greater accuracy than any model yet reported based on the Broad Bioimage Benchmark Collection [22].

**Figure 1:**
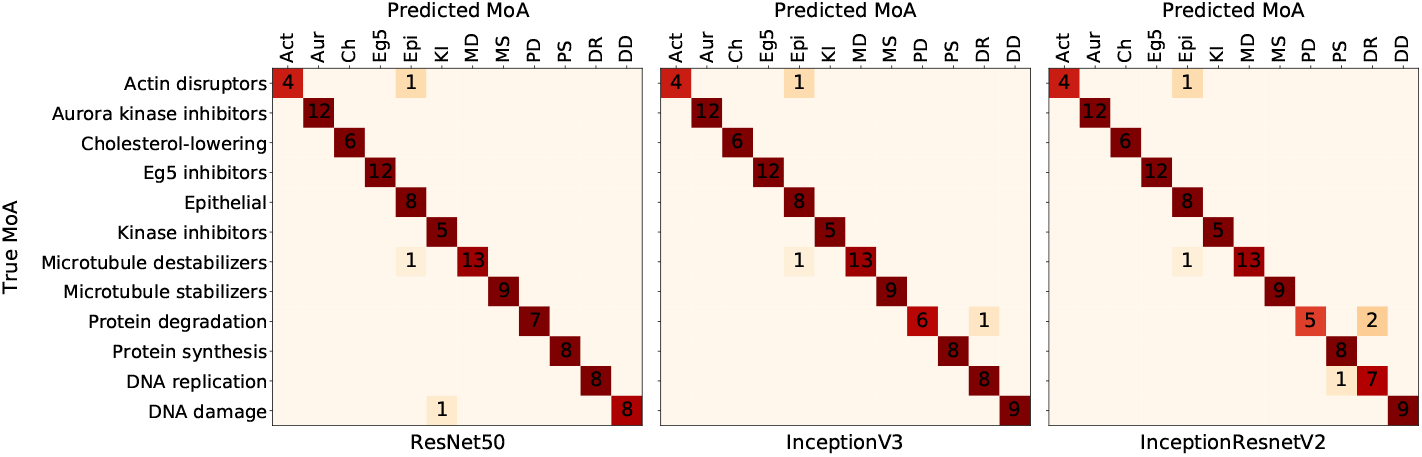
bbbc021v1 hard predictions. Confusion matrices for predictions of compound-concentration pairs, with a mean accuracy of 97%, 97% and 95% accuracy for ResNet50, InceptionV3 and InceptionResnetV2 respectively. Zeros are excluded for better visualization.

However, several of the correctly predicted compounds had a treatment soft prediction of less than 0.5 (see Additional files), which means although our model correctly predicted the MoA for these treatments, there were strong uncertainties in several of the predictions.

On the bbbc014v1 dataset the three models attained accuracies of up to 100% after just single epochs of training — an accuracy depending heavily on the stochastic process of mini-batch gradient descent. The quick learning is arguably a strong indication of transferability of the pretrained parameters. Furthermore, these results suggest that deep CNN models can be successfully applied to smaller datasets of high content imaging.

Neural networks have often been thought of as black boxes due to the difficult problem of understanding what they have learnt and how they have learnt it. For very deep neural networks, such as those studied in this paper, the problem is even more acute due to the multitude of non-linear interacting components [17]. We applied deep visualisations [17] to gain some insights into the workings of our neural networks and the transferability of the pretrained parameters (Figure 2). Notably, the early layers of the two models showed similar patterns of activation between the pre-trained and fine-tuned images, whereas the deeper layers, activated by higher level abstractions, were more dissimilar.

**Figure 2:**
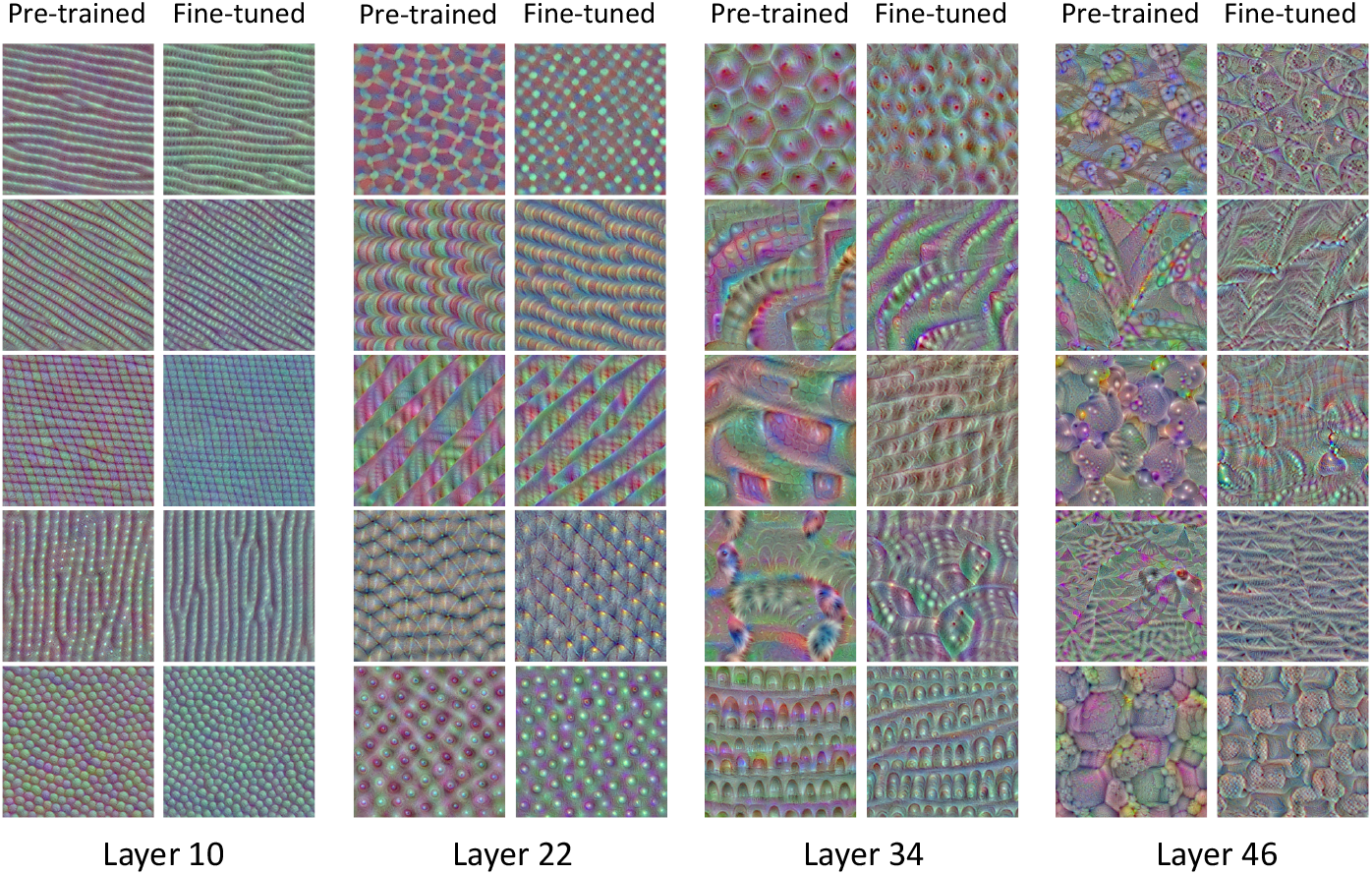
Activation maximization. A subset of images that maximize certain filter output activations in the different layers of ResNet50 using keras-vis toolkit. A comparison between the pre-trained model and the fine-tuned model.

Furthermore, the fine-tuned images of the deeper layers showed less elaborate patterns than the layers prior to fine-tuning. We speculate that this could be due to the initial networks ability to capture a greater variety of object types.

As our images are only of cells, it may be that although the pre-trained networks provide reasonable initial values for the weight parameters, for the deeper layers many of these weights will be shrunk towards zero to accommodate for this reduction in object variety.

### Conclusion and potential future work

Transfer learning and deep CNNs, when used in combination, produce highly accurate classification of MoAs. These models were able to quickly distinguish the different cell phenotypes despite a limited quantity of labelled data.

However, although the bbbc021v1 dataset is one of the very few good benchmarking datasets publicly available, it no longer presents significant challenges for many of the current state-of-the-art models, many of which have already reported accuracies of 90% and above. It would therefore be interesting to evaluate these models on more difficult classification tasks of MoAs, and evaluate them further in the field of high content imaging.

Finally, as mentioned earlier, there were strong uncertainties in many of our predictions. Formally quantifying and accounting for this uncertainty is of significant interest. An obvious avenue for this is to apply the Bayesian approach. However for deep CNNs inferring the model posterior is a difficult task and approximate methods, such as variational inference are often required [37]. A less well-known alternative approach is to utilize conformal prediction (CP). CP works atop machine learning algorithms to enable assessments of accuracy and reliability [38] and has previously been implemented with neural networks [39].

## Supplementary material

### Supplementary file 1 — GitHub Repository

This study should be entirely reproducible. Python scripts used in this study for obtaining the presented results can be found at https://github.com/pharmbio/kensert_CNN. Due to stochastic procedures, like randomly dividing the datasets into mini-batches, results will differ somewhat from session to session.

### Supplementary file 2 — Results

An Excel document containing soft predictions for each model on the bbbc021v1 dataset can be found at https://github.com/pharmbio/kensert_CNN

